# Automated Tracking of Biopolymer Growth and Network Deformation with TSOAX

**DOI:** 10.1101/316489

**Authors:** Ting Xu, Christos Langouras, Maral Adeli Koudehi, Bart E. Vos, Ning Wang, Gijsje H. Koenderink, Xiaolei Huang, Dimitrios Vavylonis

## Abstract

Studies of how individual semi-flexible biopolymers and their network assemblies change over time reveal dynamical and mechanical properties important to the understanding of their function in tissues and living cells. Automatic tracking of biopolymer networks from fluorescence microscopy time-lapse sequences facilitates such quantitative studies. We present an open source software tool that combines a global and local correspondence algorithm to track biopolymer networks in 2D and 3D, using stretching open active contours. We demonstrate its application in fully automated tracking of elongating and intersecting actin filaments, detection of loop formation and constriction of tilted contractile rings in live cells, and tracking of network deformation under shear deformation.

## 1 Introduction

Microscopic studies of biopolymer networks reveal dynamical and mechanical properties important for their function in tissues and living cells. Quantitative studies have been advanced by using *in vitro* models, where biopolymers such as actin filaments, intermediate filaments and microtubules are purified and reconstituted into networks with well-controlled conditions. Extracting quantitative information from such studies is challenging due to low image signal to noise ratio (SNR), and the large size of the data that requires automated extraction. Recently, methods and software have been developed for extraction of network structure in biopolymer images of low SNR for a single time point in 2D and 3D [Xu et al., 2015, Breuer and Nikoloski, 2015, Alioscha-Perez et al., 2016, Rogge et al., 2017, Zhang et al., 2017b]. For example, the SOAX software which is based in stretching active contours [Xu et al., 2015], uses a curvilinear network of multiple open curves (“snakes”) to automatically extract 2D and 3D networks. Very little work however exists to quantify the evolution of network structures in 2D/3D time-lapse images. Some previous efforts focused on tracking of individual filaments [Sargin et al., 2011, Kidambi et al., 2012, Smith et al., 2010] or filament tips [Applegate et al., 2011].

Automated tracking of dynamic biopolymer networks is challenging because of the variability of filament dynamics, which can include filament growth and shrinkage, filament or network node appearance and disappearance, and global or local network deformation. To study how the network remodels and deforms in time, the goal of tracking is to generate a track for each filament, segment or node in the network. Examples of studies that can benefit from automated network tracking include studies measuring the elongation kinetics of individual actin or microtubule filaments that intersect with one another as they polymerize [Fujiwara et al., 2007], measuring the deformation of cross-linked fiber networks under external forces, which relates to their mechanical properties [Lieleg et al., 2010, Sanchez et al., 2012], as well as network structures that form in cells through polymerization and depolymerization [Costigliola et al., 2017].

Formulating multi-frame object tracking as finding the minimum or maximum path cover in a *k*-partite graph has been successful for tracking cells [Xie et al., 2008], tips of microtubules [Altinok et al., 2006], and point features in natural images [Shafique and Shah, 2005]. Beside low computation complexity, the major benefit of this formulation is that it allows for false negative/positive classification of detection results (that aids when detection is influenced by image noise), and appearing/disappearing/reappearing of tracked objects. However, direct application on tracking networks of filaments does not work well because temporal consistency of network topology is not considered (see Fig. 1A). Another complication is that detection error can propagate to the correspondence phase (i.e. the process assignment of detected filaments to tracks) thus adversely affecting tracking performance. Using temporal information in the detection step has been useful in cell tracking [Schiegg et al., 2013]. The method in [Glowacki et al., 2014] improved reconstruction of tree structures by incorporating temporal information when building a spatio-temporal graph of seed points with high tubularity. No temporal correspondence was generated for individual tree branches however.

We integrated the approaches described above into a novel approach for biopolymer network tracking that (i) extends the global *k*-partite matching method to deal with curved filaments, and (ii) implements a local matching procedure during the detection phase to improve the accuracy and consistency of network topology across frames. This method has been implemented in an open source software tool, TSOAX (Trackable Stretching Open Active Contours), which is an extension of SOAX (designed for static 2D/3D images) to time-lapse movies of 2D/3D images.

**Figure 1:**
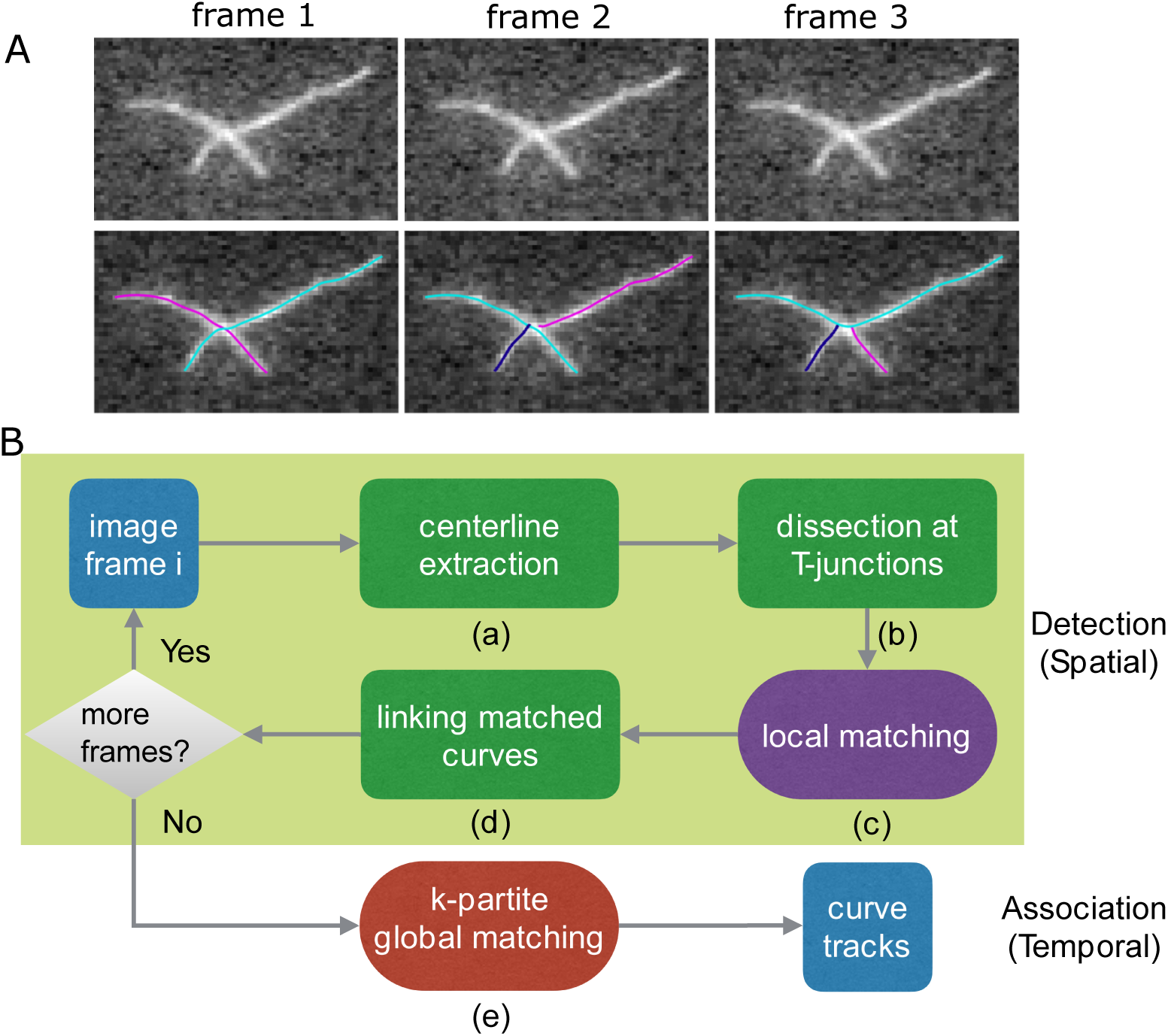
Network tracking algorithm. (A) Example of time lapse sequence (three identical frames, for simplicity). Image of actin filaments from [Fujiwara et al., 2007], 1 pixel = 0.17 *μ*m. After filament segment detection in each separate frame (shown with different colors), each frame may be extracted by a set of curves with different topology. Because of the different detected structures across frames, the network can not be tracked well using only a global matching algorithm. We thus added a local matching procedure to maintain consistent network topology. (B) Flowchart of the method consisting of a detection phase (shaded region) and a matching (i.e. temporal association) phase. The output is a set of “curve tracks,” one for each filament/segment in the network. The dark green steps are the same as in the method used in SOAX [Xu et al., 2014, Xu et al., 2015]. The steps in the other boxes are new and described in this work.

## 2 Materials and Methods

### 2.1 Outline of Computational Method

The method used in TSOAX is divided into a detection phase and a matching phase. In the detection phase corresponding to the shaded region in Fig. 1B, the centerlines of a network are extracted frame by frame using multiple open curves [Xu et al., 2014, Xu et al., 2015] (Fig. 1B(a)), which are automatically initialized and elongated sequentially. They stop elongation upon reaching filament tips or colliding with other converged curves. These curves are dissected at colliding points (“T-junctions”) which are clustered into detected network junctions (Fig. 1B(b)). This procedure uses the same method and parameter values as in the SOAX software for single frame network detection [Xu et al., 2015]. In order to produce temporally consistent network topology for the matching phase, we introduce a temporal information based local matching step (Fig. 1B(c)). Matched curves are linked together as the detection result for a filament (Fig. 1B(d)). After all the frames are processed, temporal correspondence among all the curves in the sequence are found by a global *k*-partite graph matching framework (Fig. 1B(e)).

Readers primarily interested in the software and its application to specific examples can skip directly to section 2.4

### 2.2 *k*-Partite Global Matching of Multiple Curves

Assuming extracted curves correspond to the same structures in an evolving network, we extend the *k*-partite graph matching framework to the problem of finding temporal correspondence of multiple curves in *k* frames (Fig. 1B(e)). We can construct a *k*-partite acyclic digraph *G* = (*V*, *E*) such that *V* = ⋃ _1≤*i*≤*k*_ *V_i_* where a curve *υ_i_* ∈ *V_i_* extracts filament/segment in frame *i*. A directed edge (*v_i_, v_j_*) ∈ *E* links curve *υ_i_* in frame *i* to curve *υ_j_* in frame *j* if *i < j*, i.e., an edge only points in the direction of increasing time. The edge weight *w*(*υ_i_, υ_j_*) quantifies the dissimilarity between curve *υ_i_* and *υ_j_*. A valid set of tracks is a path cover of this *k*-partite digraph, which is a set of directed paths such that each vertex *v* ∈ *V* belongs to exactly one path.

Since curves with similar position and geometry should belong to the same track, the optimal set of tracks is the path cover that minimizes the total dissimilarity of all the paths among all path covers of *G*:

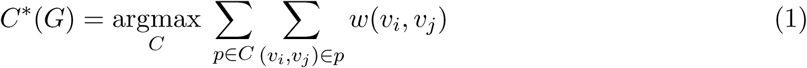

where *p* is a path in a path cover *C*.

The weight of an edge *w*(*υ_i_, υ_j_*) encodes the difference in location and geometry between curves *υ_i_* and *υ_j_*. Let *υ_i_* and *υ_j_* be represented by a linear sequence of points *R_i_* and *R_j_*, respectively, *w*(*υ_i_, υ_j_*) are computed as:

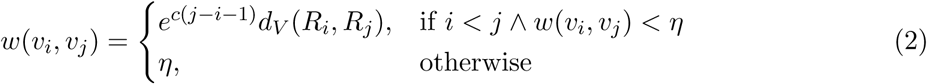

where *c* is a factor controlling how much curves from different frames are penalized and *d_V_* (*R_i_, R_j_*) is the average point distance between *R_i_* and *R_j_*,

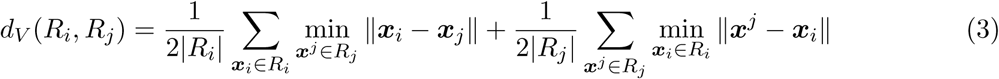

where ***x***_*i*_ and ***x***^*j*^ are points in *R_i_* and *R_j_*, respectively. A maximum weight *η* is assigned to curves that are so far apart from one another (in space or frames) that there is no need to distinguish among them. This maximum value should be at least several pixels so in the examples in this paper we set *η* = 20 pixels.

Solving Eq. 1 generates a set of tracks (paths), each containing a subsequence of curves in the original sequence. A zero-length path containing a single vertex corresponds to a singleton track containing only one curve with no predecessor or successor.

The minimum path cover can be solved in polynomial time by transforming it into bipartite matching. We construct a complete bipartite graph *B* = (*V′*, *E′*) from *G* = (*V*, *E*) as follows. The two partite vertices 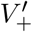 and 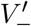 of *B* are copies of *V*, i.e. 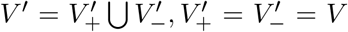. For each edge (*υ_i_, υ_j_*) ∈ *E, i < j*, we copy its weight to edge 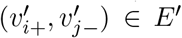. All remaining edges that correspond to matching within frames or backward matching are assigned the maximum weight *η*. It has been proved that the minimum path cover of *G* in Eq. 1 corresponds to the minimum matching of the bipartite graph *B* [Shafique and Shah, 2005].

### 2.3 Enforcing Temporal Consistency of Network Topology by Local Matching

One limitation of the above global matching is that it depends on a consistent detection result, i.e., the way extracted curves “partition” a network stays the same across frames. As in the situation shown in Fig. 1A, the accuracy of global matching will decrease when the network topology does not have temporal consistency. Therefore we introduce a novel local matching step (Fig. 1B(c)) where dissected curves are linked together respecting both local cue and temporal information from previous frames, in order to enforce a “consistent partition” of the network over time. This topological complexity is usually not present in the task of tracking point objects such as cells [Chen et al., 2015, Xie et al., 2008], tips of microtubules [Altinok et al., 2006], or other point features in natural images [Shafique and Shah, 2005].

We model this local matching as a bipartite matching as in the global matching phase. We construct a complete bipartite graph containing two copies of curve tips incident at a network junction. The edge weights are the pairwise dissimilarity 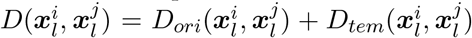 between tip 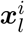 and 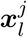 of the dissected curve 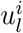 and 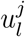 in frame *l*, respectively. The first term *D_ori_* encodes the difference of tangential orientation at tips based on the current frame,

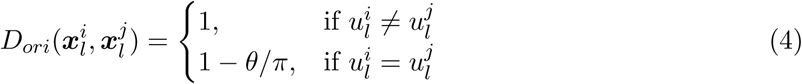

where *θ* ∈ [0, *π*) is the tangential angle between tip 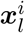 and 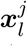. The second term *D_tem_*, utilizing temporal information, encodes the difference of the distance to the locally matched curves in the previous frame,

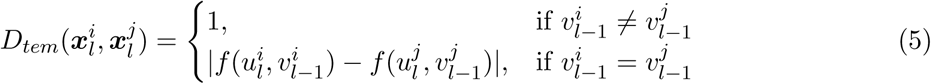

where 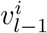 and 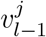 are the locally matched curve for 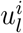 and 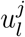 in frame *l* − 1,

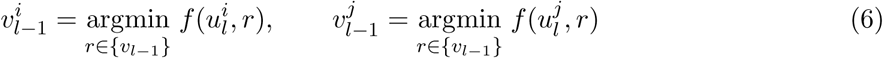

where {*υ_l−_*_1_} is the set of curves in the previous frame and 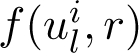 is the one-way distance between curves 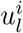 and *r*,

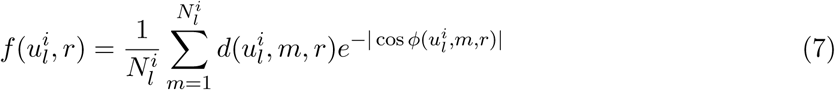

where 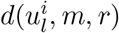 is the smallest Euclidean distance from point *m* of curve 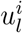 to curve *r*; 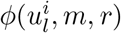 is the angle between tangent at point *m* of curve 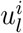 and tangent at the point of *r* that is closest to point *m*. 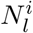 is the number of points in curve 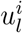.

### 2.4 Software Implementation

We implemented the above methods into the TSOAX software (http://www.cse.lehigh.edu/~idealab/tsoax), built as an extension of SOAX for static images (http://www.cse.lehigh.edu/~idealab/soax). The parameters for filament detection in each time frame are the same in both programs. Two additional parameters in TSOAX are: (1) Parameter *c* controlling the weight by which snakes detected at nearby locations in space are assigned to the same track, as a function of the number of time frames separating them (see Eq. 2). The value of 1*/c* is the number of frames beyond which the probability of assigning snakes to the same track decreases exponentially with frame number separation. This parameter should be increased (up to a value of order *c* = 1/frame) to improve track continuity over successive frames. (2) Option “Grouping,” which enables the grouping process of snakes at detected junctions prior to tracking (Fig. 2). This option can be enabled for tracking of intersecting elongating filaments (Figs. 3, 4, 5) and disabled for tracking the movement of filament segments in between junction points (Figs. 7, 8).

**Figure 2:**
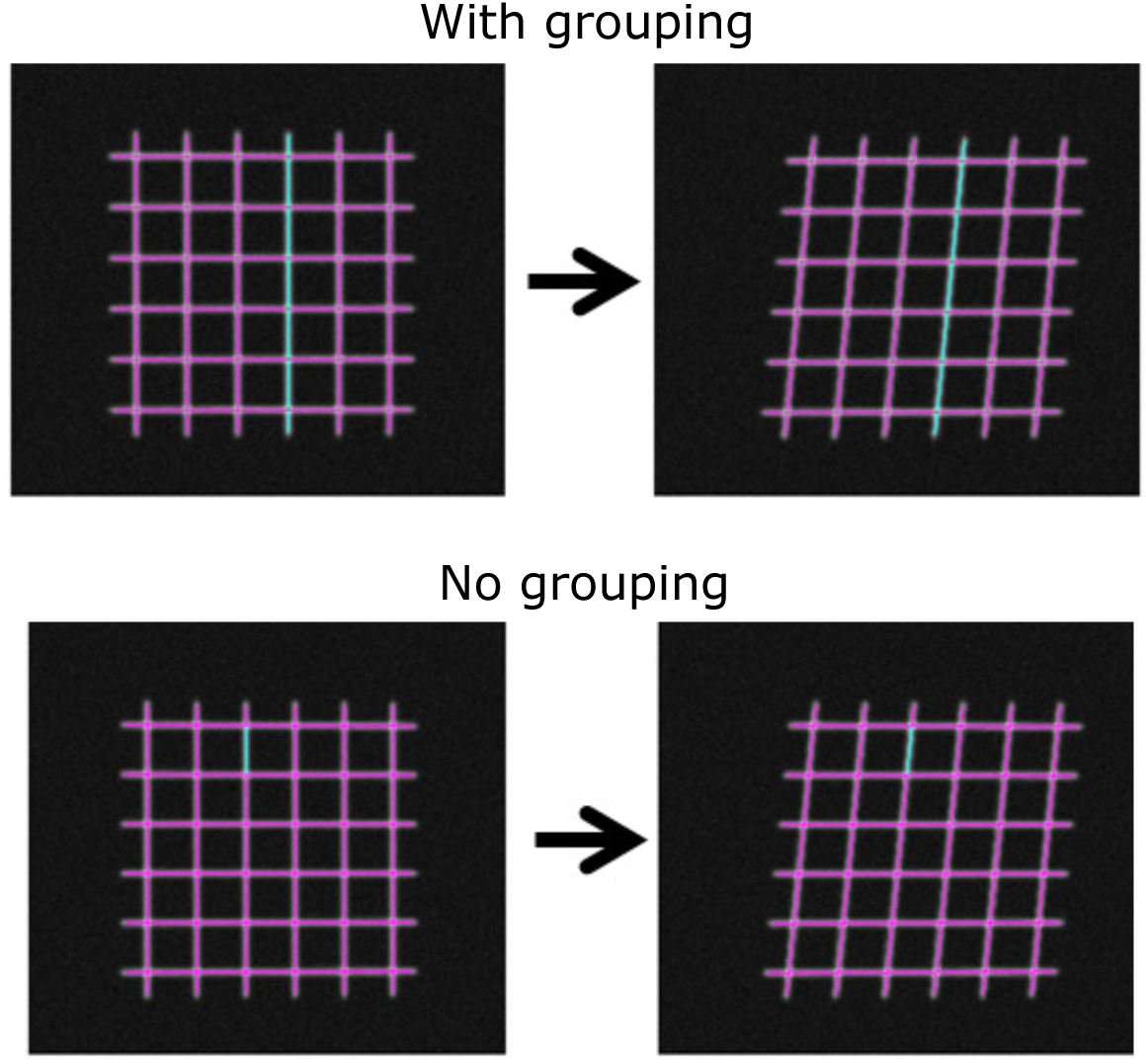
The algorithm can be applied to track fibers that extend beyond junctions (“Grouping” option in TSOAX) or just segments of fibers connecting network junction points (No grouping).

**Figure 3:**
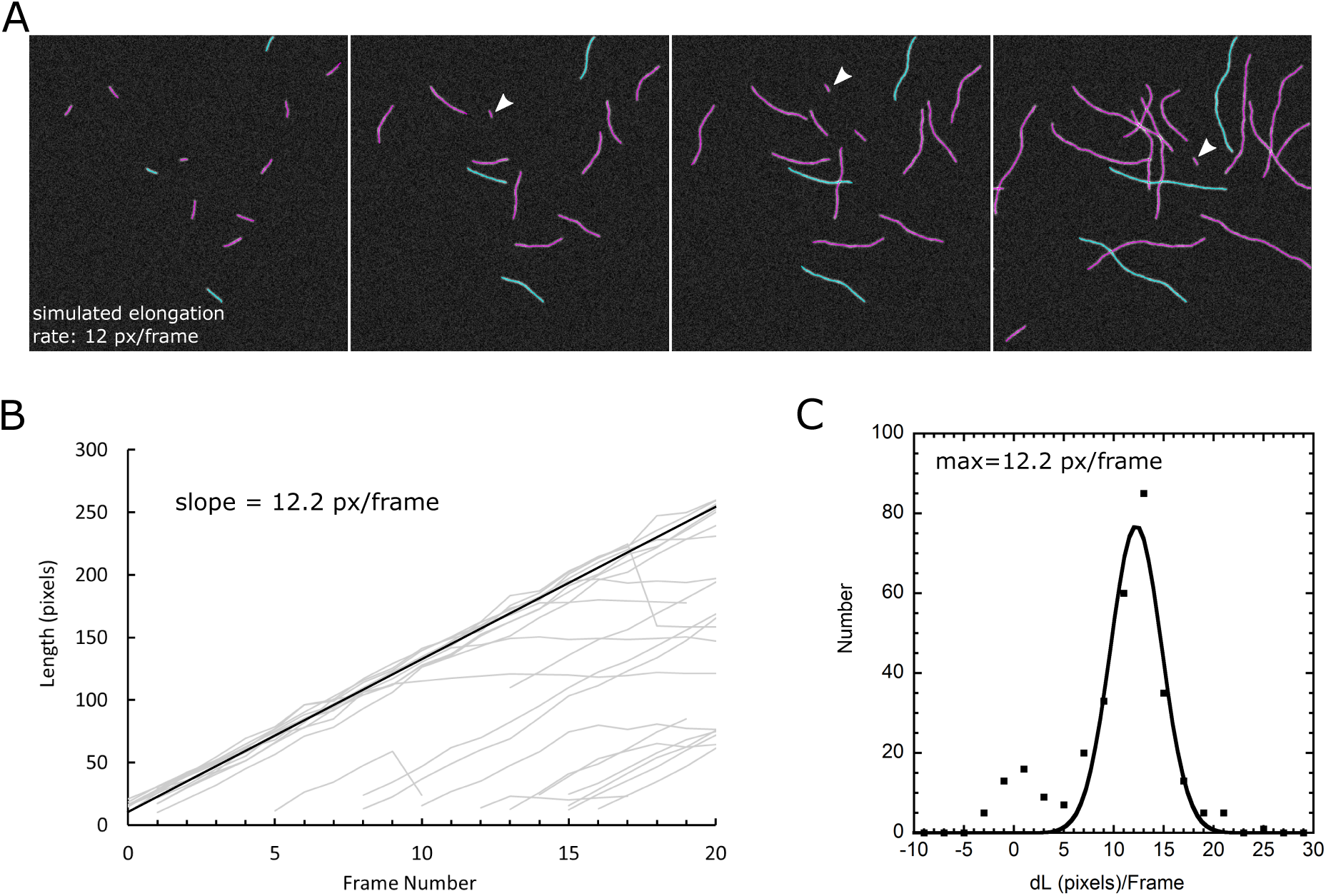
Tracking actin filaments in simulated TIRFM images. (A) Detected filaments in 2D simulated images. Arrow heads: new filaments. Blue curves show snakes belonging to the same track (three tracks shown). (B) Plot of length versus time for all tracks detected by SOAX. (C) Histogram of snake length change per frame and Gaussian fit after filtering out the data in the small peak around *dL* = 0.

**Figure 4:**
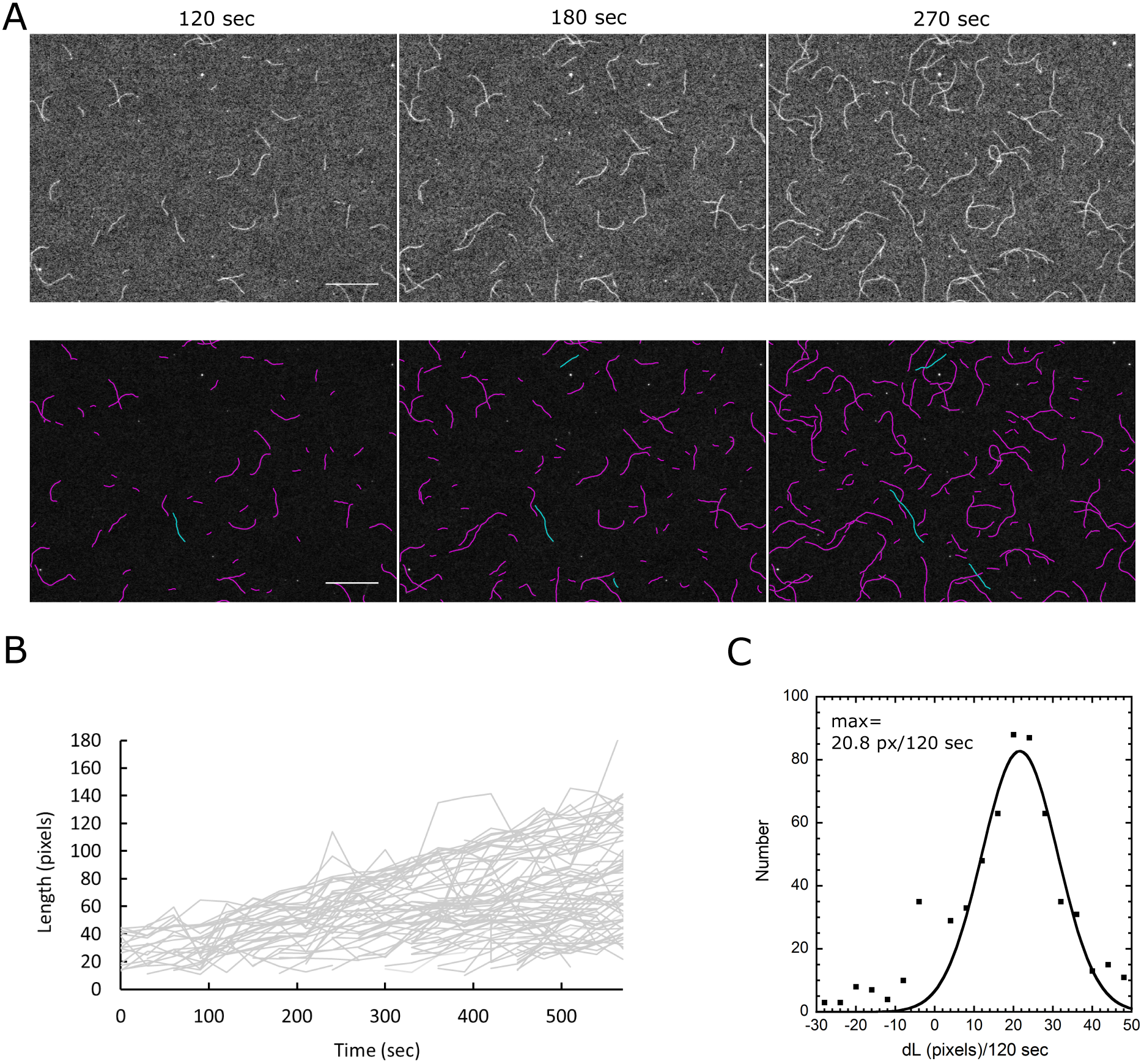
Tracking actin filaments in a TIRFM image sequence from [Fujiwara et al., 2007]. (A) First row: sample raw images. Bottom row: Tracked filaments. Blue curves show snakes belonging to the same track (three tracks shown). The pixel size is 0.17 *μ*m and the time interval between successive frames 30 sec. Bar: 20 *μ*m. (B) Plot of length versus time for all tracks detected by TSOAX. (C) Histogram of snake length change per frame and Gaussian fit after filtering out the data in the small peak around *dL* = 0.

**Figure 5:**
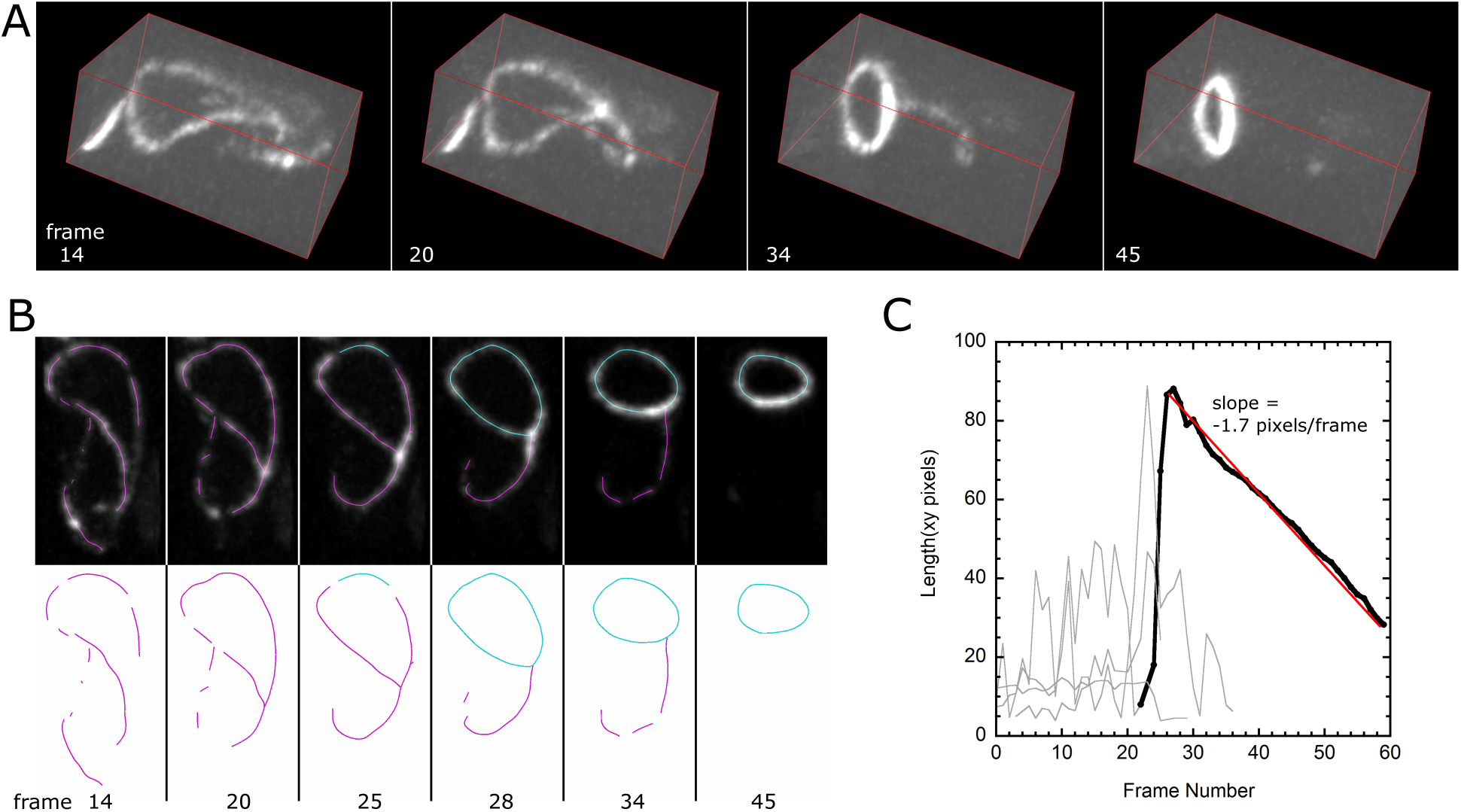
Tracking of ring formation and constriction in *mid1*∆ fission yeast cells expressing myosin light chain Rlc1-GFP. (A) Time-lapse confocal microscopy image of cells assembling a ring from Rlc1 segments distributed along the cell cortex. The cell shape is tubular and has diameter ~4 *μ*m, with its long axis along the long axis of the box. Image size: 36 × 85 pixels with 1 *μ*m = 6.95 pixels. A 3D stack of 27 *z*-slices separated by 0.3 *μ*m was obtained every 1 min. (B) Side view showing detected snakes. Highlighted in blue is the track of the the snake corresponding to the constricting ring. (C) Plot of snake segment length versus time for all tracks allows identification of the time point of ring formation and measurement of constant constriction rate 0.24 *μ*m/min (red line).

**Figure 6:**
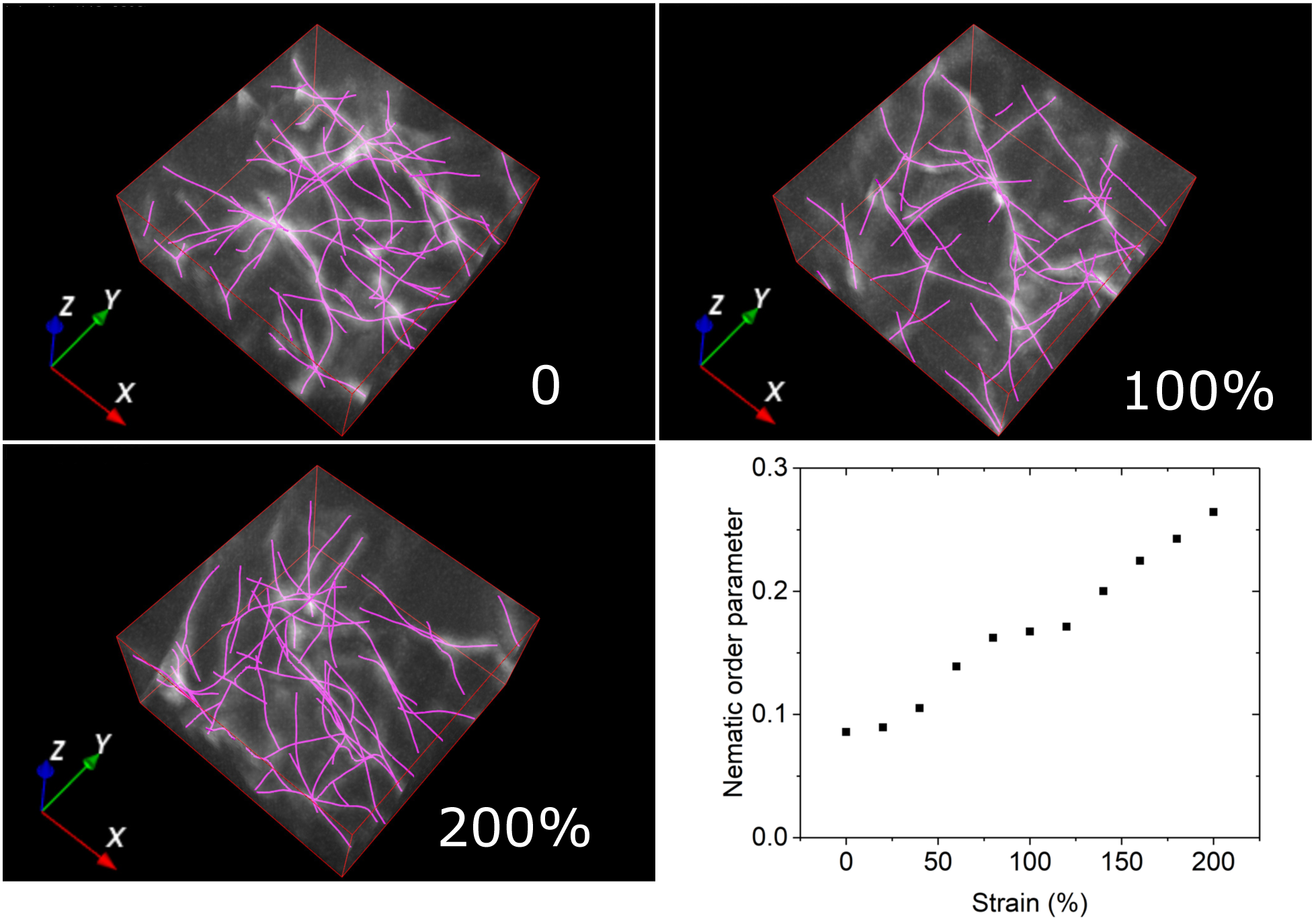
Measuring deformation of 3D fibrin networks in confocal rheology image sequence. The images show a small region of a sample of fluorescently labeled fibrin (2 mg/ml) undergoing sequential mechanical deformation along the *x* direction by an externally applied shear in 10% intervals. Images show the same region under the indicated shear and extracted network (with no grouping). The dimensions of the image along a confocal size is 16 × 16 *μ*m (100 × 100 pixels). There are 50 *z* slices, separated by 0.16 *μ*m. The graph shows the orientation-independent nematic order parameter as a function of applied strain, combining the tracked filaments from four different images (of the same size as the example in this figure) to calculate one single order parameter per strain level.

**Figure 7:**
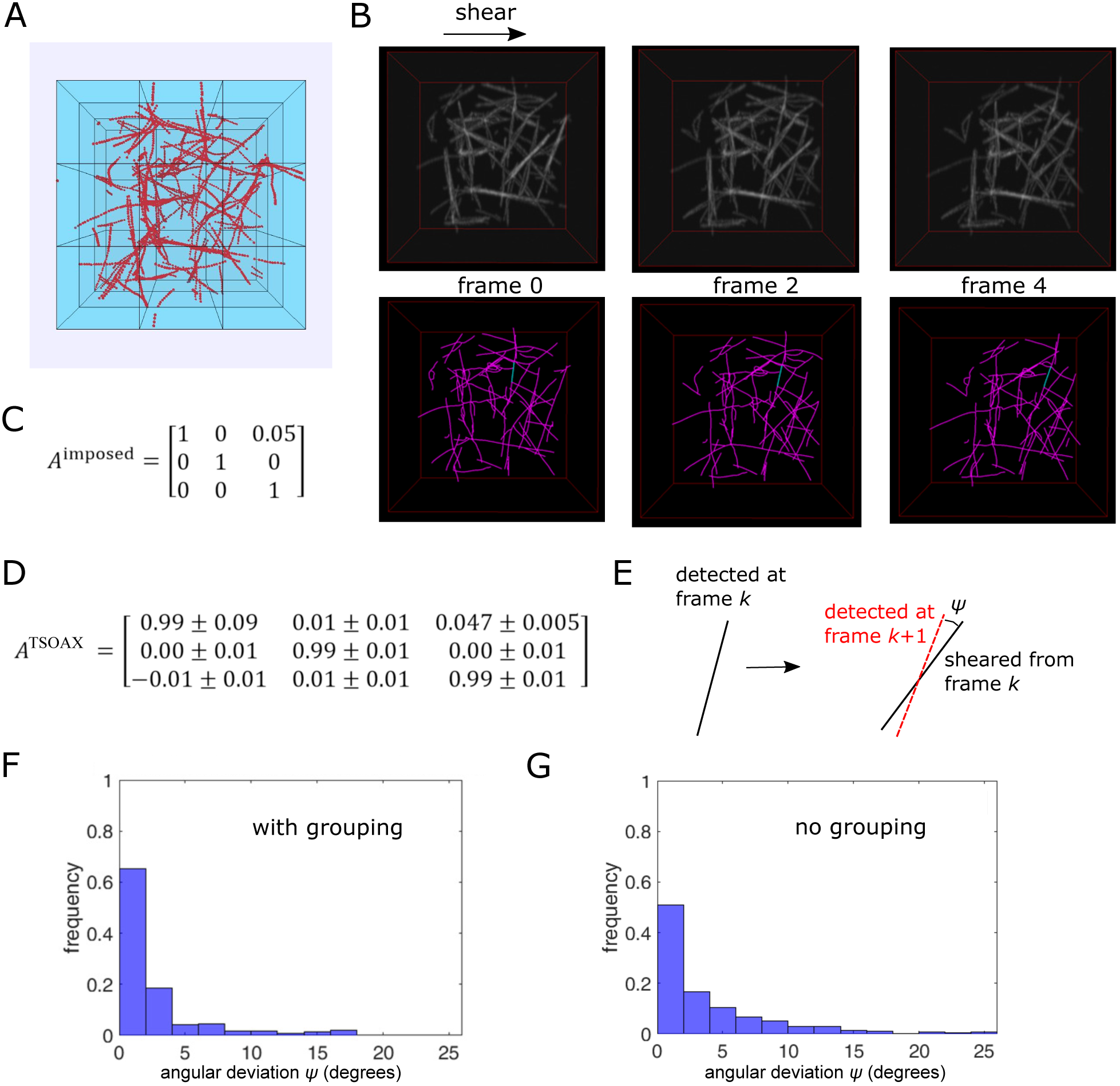
Tracking shear deformation of simulated 3D network. (A) Snapshot of Brownian dynamics simulation of actin filaments forming a network of bundles and single filaments. Simulations correspond to 3.2 *μM* actin: 120 filaments of length 1.1 *μm* in box of side 3 *μm* (whole blue box). (B) Top row: synthetic image (200 pixels/side) generated from filament coordinates in panel A under five successive shear transformations. Bottom row: TSOAX results (with no grouping) with example of tracked segment in blue. (C) Shear matrix per frame. (D) Transformation matrix calculated from TSOAX tracked snakes with grouping between frame 0 and 1 using 30 samples (10 triplets) out of 55 tracked snakes (mean *±* standard error). (E) Definition of angular deviation between axis of detected snake and axis of sheared snake of preceding frame of same snake track. (F) Distribution of angular deviation between the snakes of all successive frames of a sequence of five frames (with grouping, *n* = 237). (G) Same as F but with no grouping (*n* = 447).

**Figure 8:**
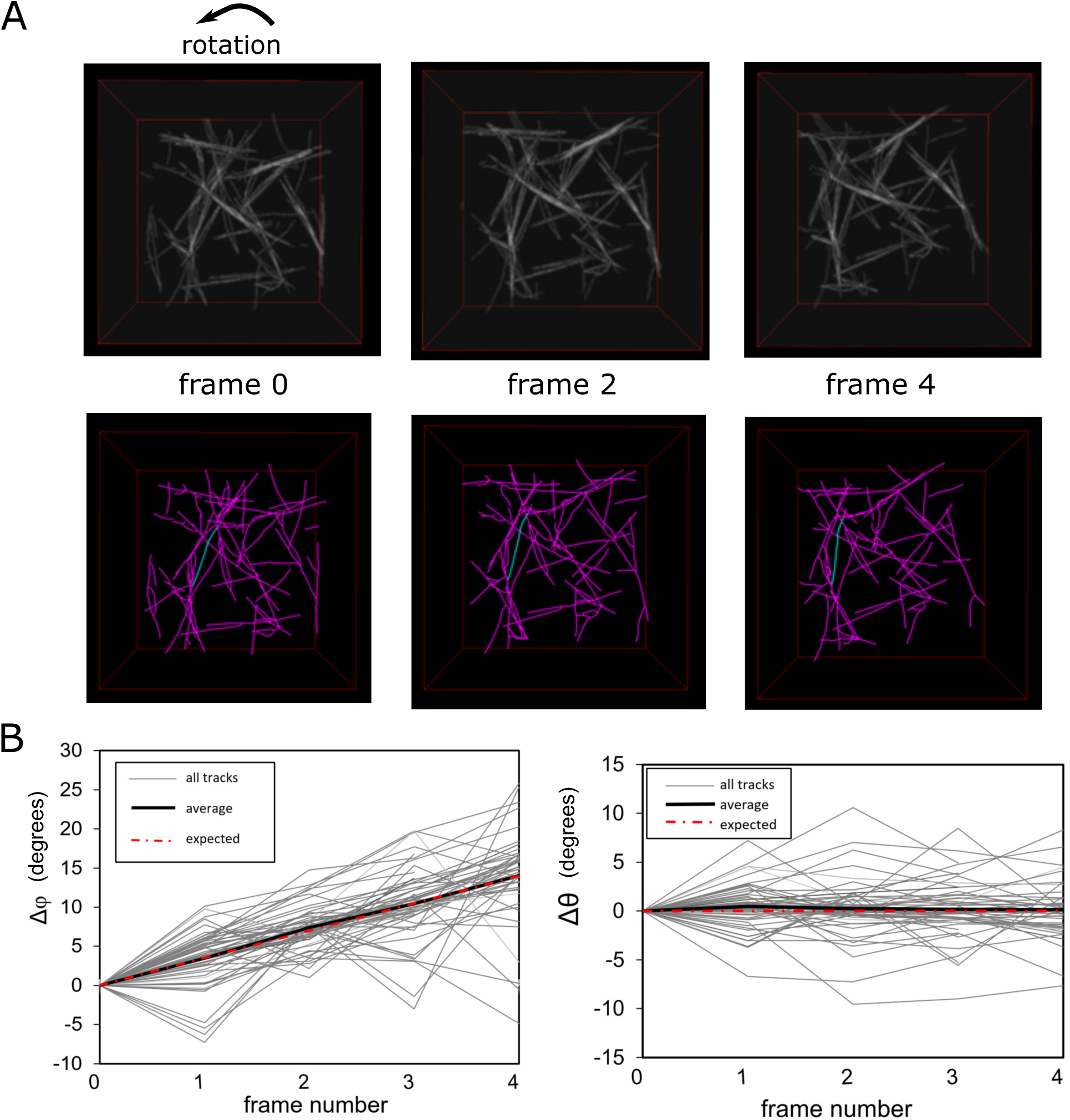
Tracking uniform rotation of simulated 3D network. (A) Top row: synthetic image (200 pixels/side) generated from filament coordinates in panel A of Fig. 7A (top view). The filaments are rotated by 3.5° along the *z* axis in a total of five iterations. Bottom row: TSOAX results (with no grouping) with example of tracked segment in blue. (C) Change of azimuthal (*ϕ*) and polar (*θ*) angle per frame vs frame number with grouping for 53 snakes presents in more than four frames.

### 2.5 Confocal Rheology of Fibrin Networks

Human plasma fibrinogen (Plasminogen, von Willebrand Factor and Fibronectin depleted) and human *α*-thrombin were obtained from Enzyme Research Laboratories (Swansea, United Kingdom). All chemicals were obtained from Sigma Aldrich (Zwijndrecht, The Netherlands). Fibrinogen was dialysed against buffer containing 20 mM HEPES and 150 mM NaCl at a pH of 7.4, and mixed with 500 mM CaCl_2_ at room temperature to a final assembly buffer containing 20 mM HEPES, 150 mM NaCl and 5 mM CaCl_2_. For fluorescence microscopy, fibrinogen labeled with Alexa Fluor 488 (Life Technologies, Bleiswijk, The Netherlands) was mixed with unlabeled fibrinogen in a 1:19 molar ratio.

Assembly was initiated by the addition and quick mixing of 0.5 U/ml of thrombin, after which the sample was quickly transferred to the confocal rheometer. Evaporation of solvent during assembly was prevented by adding a layer of low-viscosity mineral oil (M3516, Sigma Aldrich) on the liquid-air interface.

Confocal microscopy in combination with shear rheology was performed on a home-built setup, consisting of an Anton Paar rheometer head (DSR 301, Graz, Austria) placed on top of an inverted microscope equipped with a Yokogawa CSU-22 spinning disk confocal head, a Hamamatsu EM-CCD C9100 Digital Camera, and a 100x oil-immersion objective. The bottom plate consisted of a microscopy coverslip, while the top plate was a 20 mm stainless steel plate. Imaging was performed half-way the centre and the edge of the top plate. After polymerization, instantaneous strain steps of 10% were applied. Every strain step was held for 1 minute during which three-dimensional stacks were taken, starting 10 *μ*m above the glass interface. The stacks consisted of 50 steps in *z* with a spacing of 0.16 *μ* and an exposure of 0.5 s per frame.

### 2.6 Synthetic Images of Sheared and Rotated Polymer Network

We created synthetic movies of sheared and rotated polymer networks starting from a Brownian dynamics simulation of attractive semiflexible polymers (Fig. 7A). We used the method of Tang *et al.* [Tang et al., 2014] to generate polymerizing semiflexible actin filaments represented as beads connected by springs placed randomly in a 3D volume. Each filament bead represented 37 actin subunits. A short range attraction between filament beads led to the formation of a bundle network [Tang et al., 2014]. We then applied a simulated Gaussian microscope point spread function to a snapshot of the simulation and added noise to create a 3D synthetic image stack (Fig. 7B), similar to the synthetic images of Fig. S1 in Bidone *et al.* [Bidone et al., 2014]. Images of a sheared network were generated by applying successive shear transformations to the filament coordinates of the simulation snapshot, using the matrix in Fig. 7C, without applying any other movement or deformation to the filaments. For images of uniformly rotated networks we applied successive rotations of filaments by 3.5° around the *z* axis. A synthetic image was generated after each shear or rotation transformation (Fig. 7B, 8A).

## 3 Results

We apply our method to three types of biopolymer networks that indicate its broad applicability. The first is the case of polymerizing and intersecting actin filaments along a 2D slide for which we used both simulated [Smith et al., 2010] and experimental [Fujiwara et al., 2007] images. The second type is ring formation in mutant fission yeast lacking the contractile ring anchoring precursor component Mid1 [Lee and Wu, 2012]. The third type is 3D biopolymer networks undergoing shear or simple uniform rotation around an axis. For the latter case we show simulated images and an example of cross-linked fibrin bundles undergoing an externally-imposed mechanical shear deformation. Examples of these images and corresponding TSOAX parameter files can be found in the TSOAX website.

### 3.1 Tracking Actin Filaments that Intersect during Elongation

As a first test and application of our algorithm, we performed fully automated tracking and measurement of elongation rates in simulated TIRFM images (Fig. 3). In the simulated images, filaments elongated at 12 pixels per frame from one end, similar to actin filaments that elongate much faster from their barbed ends compared to the pointed ends [Pollard and Cooper, 2009]. New filaments were initiated throughout the image sequence, to simulate nucleation of new actin filaments at random location on the image. The images and added noise were constructed by modifying a method in [Smith et al., 2010].

The intersections that form in the image do not represent physical links, hence we are interested in a network of linear curves without side branches. Our algorithm takes into account the history of filament elongation, which aids in proper linking of extracted curves at junction points.

The program successfully tracked many filaments as they grew past intersection points with other filaments. The blue curves in Fig. 3A show snakes that belong to the same track (three tracks shown). A plot of length versus time for all tracks detected by SOAX is shown in Fig. 3B. Most tracks show the linear growth of 12 pixels/frame in the ground truth data, which can also be identified clearly by a histogram of the length change per frame (Fig. 3C). Tracking errors are seen as tracks with filaments that stop elongating, which correspond to failures to track a filament past an intersection (because of snakes that stopped elongating at a junction point). These events cause a smaller peak around 0 in the histogram of Fig. 3C. Other tracking errors include a sudden change in filament length and premature termination, as seen in Fig. 3B. Overall, the elongation rate was successfully extracted without the need for manual or semi-automated tracking.

We then proceeded to apply our method to experimental time-lapse sequence of fluorescently-labeled polymerizing actin filaments imaged by TIRFM (Fig. 4A). We used the data from [Fujiwara et al., 2007] for which manual tracing and semi-automated tracking with active contours of selected filaments in [Li et al., 2009] gave elongation rates of 11.2-11.3 subunits/sec. In this experiment the filaments grew parallel to a glass slide by polymerization, at higher density and noise compared to Fig. 4. As can be seen in Fig. 4B-C, the measured elongation rate with fully automated detection and tracking has a peak that corresponds to 11.0 subunits/sec after fitting the data with a Gaussian that excludes the peak of the histogram at 0. This is in good agreement with the measurements in [Li et al., 2009]. An advantage of the new method is that it includes data from all filaments on the slide, bypassing any user bias in selecting filaments for their ease for manual or semi-automated tracking.

### 3.2 Tracking Formation and Constriction of Tilted Contractile Ring

As an example of an application of our method and software in live cells, we measured the constriction rate of contractile rings in fission yeast cells. A ring can be represented as a closed snake in TSOAX. In wild type fission yeast cells the actomyosin contractile ring forms in the middle of their tubular shape, perpendicular to their long axis. However, in *mid1* ∆ cells that lack the ring anchoring protein Mid1, the ring initially assembles in a tilted and non-circular shape [Lee and Wu, 2012]. In Fig. 5A, new confocal microscopy images obtained as in [Lee and Wu, 2012] for a *mid1* ∆ cell expressing Rlc1-GFP (a light chain of type II myosin Myo2p) are shown over time. A non-circular constricting myosin closed loop forms out of linear segments associated with the cell’s cortex. The loop constricts, reorients and becomes circular over time. Our program can be used to detect and track the location of the ring precursor myosin linear elements, identify the time of continuous loop formation, and track the loop over time through the constriction process (Fig. 5B). The length versus time of tracked segments shows that the loop constricts at a nearly constant rate from the time of ring detection (Fig. 5C, red line).

### 3.3 Measuring Nematic Ordering of Strained 3D Fibrin Networks

To illustrate the application of TSOAX to 3D network images, we took time-lapse sequences of a 3D fibrin bundle network. The network was imaged by confocal microscopy with externally-imposed shear deformation in increments of 10% and up to 200% (Fig. 6). The fiber structures in the image are cables of bundled fibrin protofibrils, which establish a physically interconnected 3D network with very few free fiber ends [Piechocka et al., 2016].

Of interest in this experiment are the deformation and aligning of the fibers in response to the macroscopic external perturbation. We used TSOAX to process a sequence of images in time and calculate the change in the nematic order parameter *S* [Allen et al., 1993, Vos, 2018], a scalar metric that measures the degree of orientational order. For a completely isotropically orientated assembly of filaments *S* = 0, while *S* = 1 for fully aligned filaments. The graph in Fig. 6 shows the monotonic increase in the nematic order parameter with increasing shear strain, which signifies the alignment of fibrin fibers along the shear direction. The nematic parameter starts from a non-zero value at zero strain, which could indicate alignment due to shear forces during sample preparation as well as the minimum detectable anisotropy due to finite size fluctuations and signal to noise limitations. This analysis provides a direct way to quantify the strain-dependence of alignment, which is an important quantity because it influences the nonlinear elastic response of the network [Licup et al., 2015, Kang et al., 2009].

In Fig. 6, we focused on illustrating the ability of TSOAX to facilitate sequential network extraction, without analysis of individual fiber segment tracks that was at the limits of experimental resolution. Sequential network extraction without fiber-level tracking information is also possible in SOAX [Xu et al., 2015], however the workflow is more much more involved as it requires setting up a batch process over a directory of separate 3D images.

### 3.4 Tracking Simulated 3D Polymer Network Undergoing Shear or Rotation

Mechanical properties of biopolymer networks depend on the deformation and dynamics of individual network segments [Zagar et al., 2015]. To further demonstrate the use of TSOAX in tracking network deformation and dynamics, we created a synthetic movie of a polymer network undegoing shear deformation starting from a Brownian dynamics simulation of attractive semiflexible polymers (Fig. 7A-C). TSOAX was applied without (Fig. 7B) or with snake grouping. The program was able to correctly track the main features of the network. We were able to accurately recover the transformation matrix between the first two frames: the nine elements of matrix *A* in Fig. 7C were determined by selecting three snakes tracked between the first two frames, finding the unit vector along each snake direction, and then solving for the transformation of the three unit vectors of frame 0 to the three unit vectors of frame 1. We averaged the elements of *A* over 10 such triplets.

Non-affine deformations of sheared polymer networks are predicted to be important in determining their mechanical behavior [Zagar et al., 2015]. To show how accurately TSOAX can capture the affine transformation of Fig. 7B, we measured the angle between the axis of a snake detected at frame *k* + 1 and the snake of frame *k* of the same snake track after applying the imposed shear transformation (Fig. 7E). This angle would be zero in the limit of perfect detection and tracking. We found that most such angles are less than 5° (Fig. 7F,G). This illustrates that non-affine deformations larger than this angle may be detectable by TSOAX (for the conditions of this example). The few events with large angles in the histogram of Fig. 7F,G correspond to tracking errors, typically as a result of differences in topology and junction detection between successive frames. These errors were less frequent with grouping turned on (Fig. 7F,G).

We also applied successive rotations of filaments by 3.5° around the *z* axis and tracked the rotated network with TSOAX (Fig. 8A). For this transformation, the azimuthal angle of a line along each snake segment *ϕ* (measured as in [Xu et al., 2015]) should increase by 3.5° per frame while the polar angle *θ* should remain the same. The change of *ϕ* and *θ* of the detected snake tracks are shown in Fig. 8B. The average value agrees well with the imposed rotation, to within fluctuations due to segmentation and tracking errors.

## 4 Discussion

Application of our network tracking method to experimental images shows its use in a big range of 2D and 3D networks. We showed example of sequences up to 60 frames, containing 1-200 converged snakes per frame. For such image stacks, the time required for TSOAX to establish snake correspondence is typically less than the snake detection time (though the latter can become limitting for more snakes per frame and more frames). For large images, improvements in snake convergence speed may be achieved by using a smaller density of snake points (such as 1 point every 3 pixels) and reducing the minimum required number of iterations per snake evolution (e.g. reducing parameter “Maximum iterations” to 100). Even without the tracking functionality, the program is useful in network detection over multiple time frames (Fig. 6). The data of filament and junction coordinates and tracking correspondence are saved as a text file to enable analysis depending on the particular application.

The accuracy of the tracking results is limited by the accuracy with which the network of snakes is extracted in every frame. It was shown in [Xu et al., 2015] that a value of a signal to noise ratio (defined in the local neighborhood of snake point [Xu et al., 2015]) of order 5 is needed for reliable automated extraction in SOAX, after using a procedure to optimize the parameters based on an optimization function. Since TSOAX uses the SOAX alogorithm for network extraction for each time frame, the same signal to noise ratio limitation and procedure to optimize parameters also applies in TSOAX. We described how tracking snakes in time introduces additional complexities that may limit track accuracy in ways that vary depending on the particular application. We illustrated different ways by which the accuracy of the network tracking can be evaluated. This included simulated time-lapse images with added noise for the case of actin filament elongation. For deforming networks, we imposed a known affine transformation to an initial image generated by a simulation. A similar procedure of applying a transformation to an initial experimental image can be used to evaluate tracking accuracy.

Software developed to track gliding filaments for in vitro motility assays used skeletonization to extract linear filaments in each frame [Aksel et al., 2015]. Tracks of gliding filaments were constructed by connecting filaments between successive frames based on distance criteria (similar to Eq. 2) in addition to criteria that favor gliding along the longitudinal direction (not included in TSOAX). Crossing filaments that occur during gliding filament collisions were excluded from the analysis. TSOAX may be applicable to such systems while also able to track filaments during collisions.

The method and associated TSOAX software should be applicable to a broad class of systems beyond those presented in this paper. This includes images of mitochondrial networks (for which software exists for static images [Viana et al., 2015]), images of amyloid and cellulose fibers [Picu and Sengab, 2018, VandenAkker et al., 2016], plant cell wall deformation imaged by by AFM [Zhang et al., 2017a], and images of carbon nanotube networks [Bedewy et al., 2009]. It may also form a basis in future work to track more complex events such as filament merging, splitting, and overlapping.

## Author Contributions

TX developed TSOAX methods and software in collaboration with XH and DV and feedback from all authors. CL and DV performed analysis and testing for Figs. 3, 4 and 5. NW performed experiments for Fig. 5. BEV and GHK performed experiments and analysis for Fig. 6. MAK generated images and performed analysis for Fig. 7 and 8 with input from DV. TX, XH and DV wrote the paper with input from all authors.

## Acknowledgements

This work was supported by NIH grants R01GM114201 and R01GM098430. GHK and BEV acknowledge support from the the Netherlands Organisation for Scientific Research (NWO) and the Foundation for Fundamental Research on Matter (FOM Program grant nr 143). We thank Jian-Qiu Wu for his help with the experiments with fission yeast.

